# From foldamers to functional pores: a force field for oligourea-based desalination channels

**DOI:** 10.64898/2026.04.21.719837

**Authors:** Julie Ledoux, Fabio Sterpone, Marc Baaden

## Abstract

Sea water desalination is a critical solution to global water scarcity, primarily relying on membrane-based technologies and reverse osmosis. Artificial water channels (AWCs) offer a promising solution for next-generation desalination membranes. Very promising channel building blocks for AWC is oligourea foldamers repeats. These biomimetic molecules, composed of unnatural amino acids, self-assemble into protein-like superhelical channels with water-filled pores, making them ideal candidates for selective water transport. *In silico* studies provide essential support to experimental efforts in designing novel oligoureas. However, such studies require a dedicated force field for now unavailable for these molecules. In this work, we developed a tailored force field for oligoureas by adapting parameters from two established protein force field families: CHARMM (CHARMM36m) and Amber (GAFF2), using their structural similarities to natural amino acids. Our objective was to identify a force field that reliably preserves the structural integrity of oligourea foldamers. We evaluated two distinct oligoureas using molecular dynamics (MD) simulations across increasing system sizes. By comparing simulation results with experimental data, we assessed key structural features, including folding patterns, stability, and pore shapes. Our findings demonstrate that the CHARMM-based force field consistently reproduces experimental observations for both foldamers, outperforming the Amber-based alternative. This newly developed CHARMM-based force field paves the way for further exploration of oligoureas, enabling deeper insights into their stability and efficiency in sea water desalination.

## 1 Introduction

The global freshwater crisis represents one of the biggest challenges of the 21st century, with half of the world’s population currently affected by water scarcity [1]. This growing issue is driven by many causes such as population expansion, industrialization, deforestation, and the accelerating impacts of climate change that disturb the natural water cycle [2, 3]. In this context, desalination has emerged as one of the main technological solutions, offering a means to convert seawater into freshwater that is safe for drinking or use in agriculture [4]. At the forefront of this effort are membrane-based technologies [5]. However, such membranes come with significant drawbacks, particularly their energy requirements and the high costs associated with design, maintenance and reuse, particularly under constraining conditions such as in the reverse osmosis (RO) process [6]. These limitations have created a demand for alternative approaches that can offer better efficiency and sustainability.

Among the many membrane technologies proposed, biological inspirations emerge as encouraging concepts. In particular, aquaporins (AQPs), a class of transmembrane proteins that facilitate the fast transport of water molecules across cell membranes [7], are promising starting points for new membrane-based technologies for water desalination. Despite their efficiency, practical usage of AQPs suffers from critical issues: they are unstable under the high-pressure and high-salinity conditions required for industrial desalination with RO processes [8]. This instability has motivated the development of artificial water channels (AWCs). Synthetically designed AWCs aim to replicate the functionality of AQPs while bypassing their limitations.

Artificial water channels represent a new approach to desalination, combining the potential of biological systems with the advantages of engineered materials [9, 10]. Peptide-based systems, though advantageous in terms of modularity, are constrained by the relationship between their primary sequence and secondary structure [11]. This limitation has led researchers to explore foldamers, synthetic oligomers designed to mimic the folding behaviors of natural proteins [12–15].

A significant advancement has been achieved with oligoureas, a class of non-peptidic foldamers that enable precise control over self-assembly in aqueous environments [10]. Structurally, oligoureas differ from natural amino acids in their backbone composition: natural amino acids feature a backbone with the central NH–CHR–CO motif, and oligoureas possess an extended backbone NH–CHR–CH_2_–NH–CO (where R represents the side chain). This modification confers unique properties, enhancing their potential for controlled self-assembly. Recent research has demonstrated that oligoureas can self-assemble into a variety of protein-like quaternary structures, ranging from discrete helical bundles to dynamic superhelical channels featuring water-filled pores [10].

To further accelerate the design and optimization of synthetic molecular systems, computational modeling has become a powerful tool to complement experimental approaches. Among these techniques, molecular dynamics (MD) simulations stand out for investigating the structural and dynamical properties of biomolecular systems, including proteins, nucleic acids, lipids, and small molecules. MD simulations rest on Newton’s equations of motion to model the time-dependent behavior of molecular systems under the influence of interatomic forces. Central to this approach is the molecular potential energy function necessary for the integration of Newton’s equations. Collectively known as a force field, it requires a comprehensive set of parameters to define the interactions between all atoms in the simulation. The simplest molecular potential function integrates both bonded and non-bonded interaction terms. Bond stretching and angle bending are typically described using harmonic potentials, while dihedral angles are defined by periodic functions. Non-bonded interactions contain the van der Waals forces, modeled via a Lennard-Jones potential, and electrostatic interactions, represented by a Coulomb potential.

Oligourea residues share structural and chemical similarities with natural and unnatural amino acids, differing primarily in their backbone composition (Figure 1A). Despite the availability of force fields for various unnatural amino acids across different force field families [16–20], and the extensive efforts of the scientific community in developing machine-learned force field parameters for a wide range of molecules, there are currently no oligourea-type molecule parameterizations offered to the scientific community. Given the absence of accessible parameters in the literature, we decided to parametrize our own force field, specific to oligourea foldamers with amino-acid side-chains.

**Figure 1:**
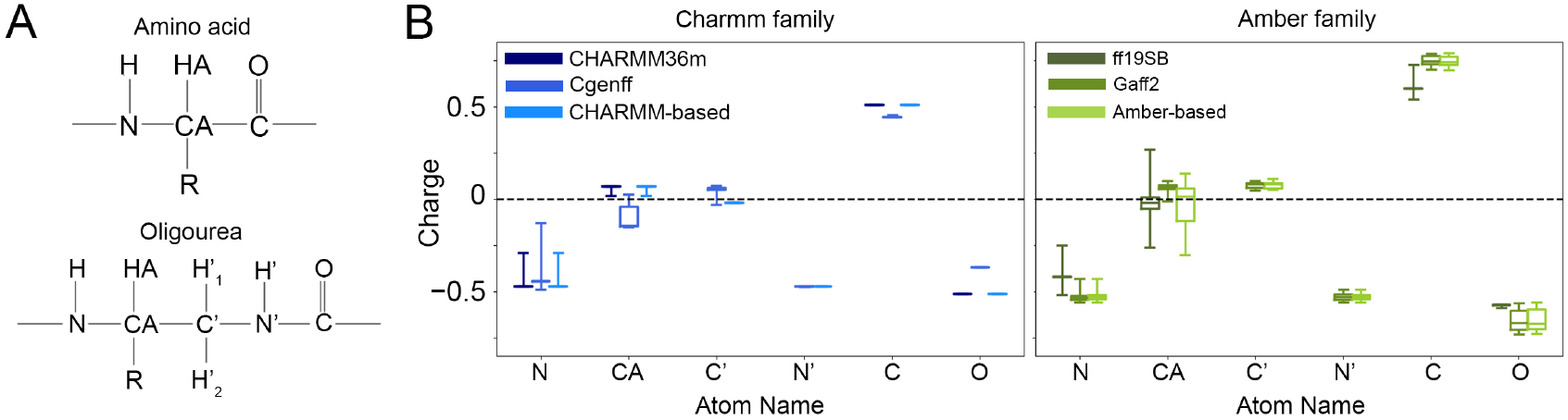
A. Oligourea general formula (R: a canonical amino acid sidechain). B. Distribution and comparison of the assigned partial charges of the backbone of the 20 “canonical” oligoureas with the partial charges of the canonicals amino-acids as parametrized in CHARMM36m, CGenFF and ff19SB force fields. C’ and N’ atoms only have values with GAFF2 and the Amber-based force fields as these atoms are not part of a canonical amino-acid in the CHARMM and Amber families force fields. Hydrogen atoms are not shown.

We chose to use two established non-polarizable biomolecular protein force fields as a starting point for developing an oligourea-specific force field: CHARMM36m (CHARMM)[21] and ff19SB (Amber)[22], taking advantage of their similarities with proteins. Their ability in reproducing the structural and dynamical features of two representative oligourea foldamers will be benchmarked against their available experimental data.

Ultimately, our goal is to identify the most suitable force field among the two parameterizations for future *in silico* studies to assist in the design of oligourea foldamers for applications in water desalination. Our tools will facilitate the setup of any novel oligourea foldamer sequence for molecular dynamics simulations.

## 2 Results

### 2.1 Partial charges and atom types assignation

In molecular force fields, atoms are characterized by their mass, partial charge, and parameters describing non-bonded interactions. All features are encoded within their atom type. Each atom type defines a specific set of parameters for both bonded and non-bonded interactions between two or more atoms. Alongside partial charges, atom types are the critical components that must be assigned when developing a new force field based on and derived from existing biomolecular force fields.

CHARMM and Amber force fields differ significantly in their approach to partial charge parameterization used to compute the pair electrostatic interactions. CHARMM uses condensed-phase interaction energies with water, whereas Amber adopts a gas-phase quantum mechanical (QM) electrostatic approach.

To transition from amino acids to oligoureas, the atoms C’ and N’, along with their respective hydrogens, are introduced into the backbone (Figure 1A). C’ is an aliphatic carbon bonded to one of the two nitrogens of a urea group. The closest small-molecule analogs to this functional group (urea and 1,3-dimethylurea) are already parametrized in CGenFF (CHARMM General Force Field) [19] and in GAFF2 (General Amber Force Field) [23, 24]. A comparison of the partial charges assigned to the two nitrogens in both urea and 1,3-dimethylurea reveals that the charges are identical between the respective nitrogens within each force field (Table S1, S2).

#### 2.1.1 Charmm-based force field

In the protein force field CHARMM36m, the partial charge assigned to the backbone nitrogen atoms of amino acids is -0.47 (with the exception of proline), while the charge for the bonded hydrogen is +0.31. Based on our observations of simple urea-containing molecules, we assigned the partial charges of N’ and H’ to match those of the NH group, using the same atom types.

For the CH_2_ group, the atom types for C’ and H’ were defined as those of an aliphatic CH_2_. The partial charge for H’ was set to +0.09, consistent with an aliphatic hydrogen. The partial charge for C’ was then adjusted to ensure the net charge of the oligourea matched the desired value (0, +1, or 1), depending on the type of side chain (Figure 1B).

Since the side chains of oligoureas are identical to those of amino acids, we directly transferred the corresponding parameters from CHARMM36m to our oligourea-specific force field variant.

#### 2.1.2 Amber-based force field

To maintain consistency with the Amber parametrization scheme, we opted to use the GAFF2 force field with AM1-BCC charges. The AM1-BCC charge model was developed as an alternative to gas-phase quantum mechanical (QM) charge fitting, which was employed for the ff19SB force field. However, we observed that the total charge of each oligourea did not align with the desired net charge (0, +1, or 1).

When comparing the assigned charges with those of ff19SB, we observed a discrepancy in the charge distribution at the C*α* atom which we corrected. Specifically, we adjusted the charge of the C*α* atoms to ensure that each oligourea residue achieved its target total charge. Following this correction, the C*α* charge distribution in our Amber-based force field closely aligns with that of ff19SB (Figure 1C).

A notable distinction exists between CHARMM- and Amber-based force fields: the Amber-based force field exhibits an enhanced electrostatic character in the urea group. This suggests that interactions within the backbone or with water may be marginally weaker in the CHARMM-based force field, potentially influencing the structural and dynamical properties of the simulated foldamers.

### 2.2 Oligourea helices

To evaluate the performance of our CHARMM- and Amber-based force fields in modeling oligoureas, we conducted MD simulations to explore the conformational ensembles of two model oligourea foldamers and compared them against data obtained from experiment. These foldamers were selected for their similar length but differing sequences, physicochemical properties, and assembly behaviors.

Whatever their sequence, oligourea foldamers tend to adopt a canonical 2.5-helix conformation [25], stabilized by a hydrogen-bonding pattern involving the oxygen of residue *i*, the N nitrogen of residue *i−* 2, and the nitrogen of residue *i−* 3. This secondary structure is characterized by three dihedral angles: Φ (C–N–C*α*–C’, average: –103°), Θ_1_ (N–C*α*–C’–N’, average: +57°), and Θ_2_ (C*α*–C’–N’–C, average: +80°). Another interesting property to examine are the B-factors obtained from crystallography that reflect the temperature-dependent atomic vibrations and are proportional to the root mean square fluctuation (RMSF) of atomic positions [26] in the crystal. As oligourea foldamers self-assemble to form structured pores [27], we assessed structural features not only at the level of individual helices but also for crystal patches of increasing size, designed to closely mimic the crystallographic data. For these systems, the pore shape and size served as an additional comparative feature. The three-dimensional structures of both foldamers were previously experimentally determined using X-ray crystallography and are available in the *Cambridge Crystallographic Data Centre* (CCDC). To ensure consistency with experimental data, our *in silico* studies retained the chemical protecting groups capping the foldamers. For clarity, oligourea residues will hereafter be denoted as uX, where X corresponds to the one-letter code of the analogous canonical amino acid.

#### 2.2.1 Aliphatic C5 helix

The C5 helix (CCDC: 836812) is a 13-oligourea aliphatic foldamer with the sequence Boc-[uV-uA-uL]_2_-uP-[uV-uA-uL]_2_-NMe (where Boc is a tert-butoxycarbonyl and NMe is a methylamide group) [28]. While this helix is not directly relevant for water translocation in future membrane applications, it provides a useful benchmark for testing our two force fields.

Due to its aliphatic nature, the C5 foldamer was investigated in both water and methanol solutions. Since the results were consistent across solvents, we focus here on the results obtained in aqueous solution.

After 100 ns MD simulation, the initial helix undergoes partial to significant unfolding depending on the force field (Figure 2A). Notably, the system simulated with the CHARMM-based force field demonstrates a tendency to maintain helical folding along the trajectory, suggesting a potential stabilization effect not observed with the Amber-based force field (Figure 2B). The helical content was monitored over time for both force fields. With the CHARMM-based force field, the foldamer maintains an average of 10 helical residues (out of 15). In contrast, with the Amber-based force field, the helix shows a gradual decrease in helicity after which only 1 to 3 residues remain helical.

**Figure 2:**
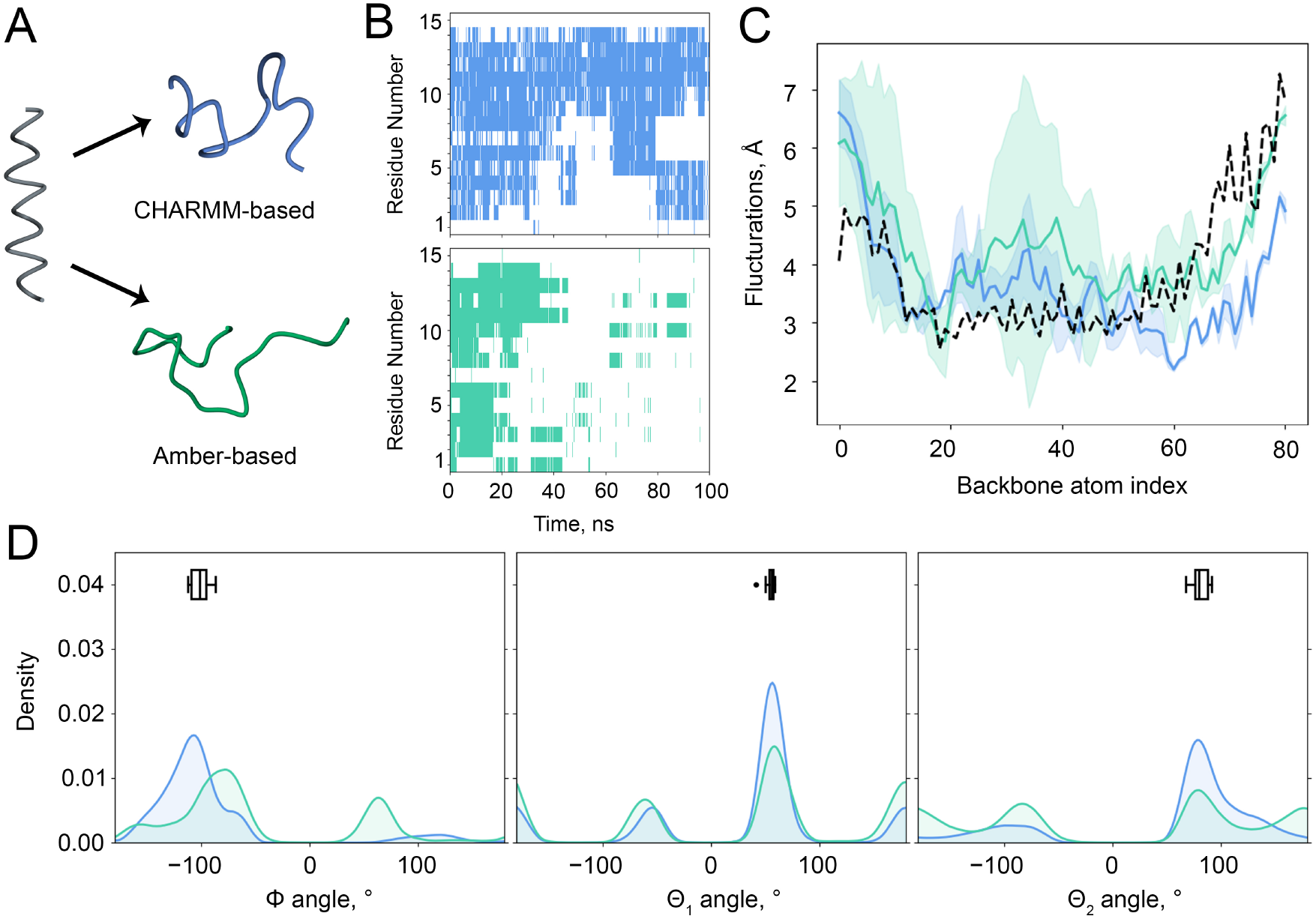
A. Last conformations sampled from the crystallographic structure of C5 for both CHARMM-(blue) and Amber-based force fields (green). B. Time evolution of the helicity by residue. C. Backbone RMS fluctuations compared to the experimental B-factors (bold dashed line). D. Distribution of the backbone dihedrals Φ, Θ_1_ and Θ_2_. For all figures, the crystallographic structure is represented in black (dashed lines, crosses), CHARMM-based force fields in blue, and Amber-based force field in green.

The root mean square fluctuations (RMSF) of backbone atoms were compared to the crystallographic B-factors (Figure 2C). With both force fields, the systems show greater fluctuations than in the experimental data, particularly between uL4-uA10 and uL4-uV12. Despite overall higher fluctuations, the overall patterns can be compared, indicating that the system simulated with the CHARMM-based force field aligns more closely with the crystallographic B-factors than for the Amber-based one. Other important structural features are the three key backbone dihedral angles. Their distributions have several maxima compared to a single reported crystallographic value. They show Gaussian mixture distributions with the crystallographic value corresponding to the maximum.

For the Φ angle, the helix simulated using the CHARMM-based force field shows a single peak closely matching the experimental average, whereas with the Amber-based force field, the torsion exhibits two peaks, deviating significantly from the expected values (Figure 2D, left). On Θ_1_, with both force fields, the distribution display three peaks at similar angles. However, with the CHARMM-based force field, the simulation samples the experimental values more effectively (Figure 2D, middle). Looking at Θ_2_, both force fields sample in the experimental torsions with an additional sampling on the opposite angle. Despite theses broader sampled regions, the CHARMM-based force field more often reflects the experimental values (Figure 2D, right).

These results suggest that the CHARMM-based force field has a tendency to better preserve helicity and reproduce experimental torsion angles. This observation holds despite the Amber-based one featuring more pronounced electrostatic characterization of the urea group. The fact that the enhanced electrostatic interactions in the Amber-based force field did not translate to improved helical stability may indicate that the dihedral parameters play a more critical role in maintaining folding. However, we cannot judge whether the preservation of helicity is to be expected or not, as we lack experimental data for the aqueous solution phase. Therefore, this metric marks a clear distinction in both force field parameterizations, but does not represent a selection criterium for deciding which one reproduces the intrinsic properties more accurately.

#### 2.2.2 Amphiphilic H5 helix

To further evaluate their performance, we extended our analysis to a second amphiphilic oligourea foldamer.

Helix H5 (CCDC: 1030454) is a promising hydrosoluble 12-oligourea foldamer with the sequence iPr-uA-uL-uK-uL-uE-uY-uL-uE-uL-uK-uA-uL-NH_2_. This foldamer self-assembles in solution to form a nanopore structure, making it highly relevant for applications in water filtration [27]. The crystallographic unit of H5 consists of a homodimer of helices, stabilized by salt bridges between the uK and uE residues (Figure 3A). Due to its amphiphilic nature, H5 was studied exclusively in aqueous solution.

**Figure 3:**
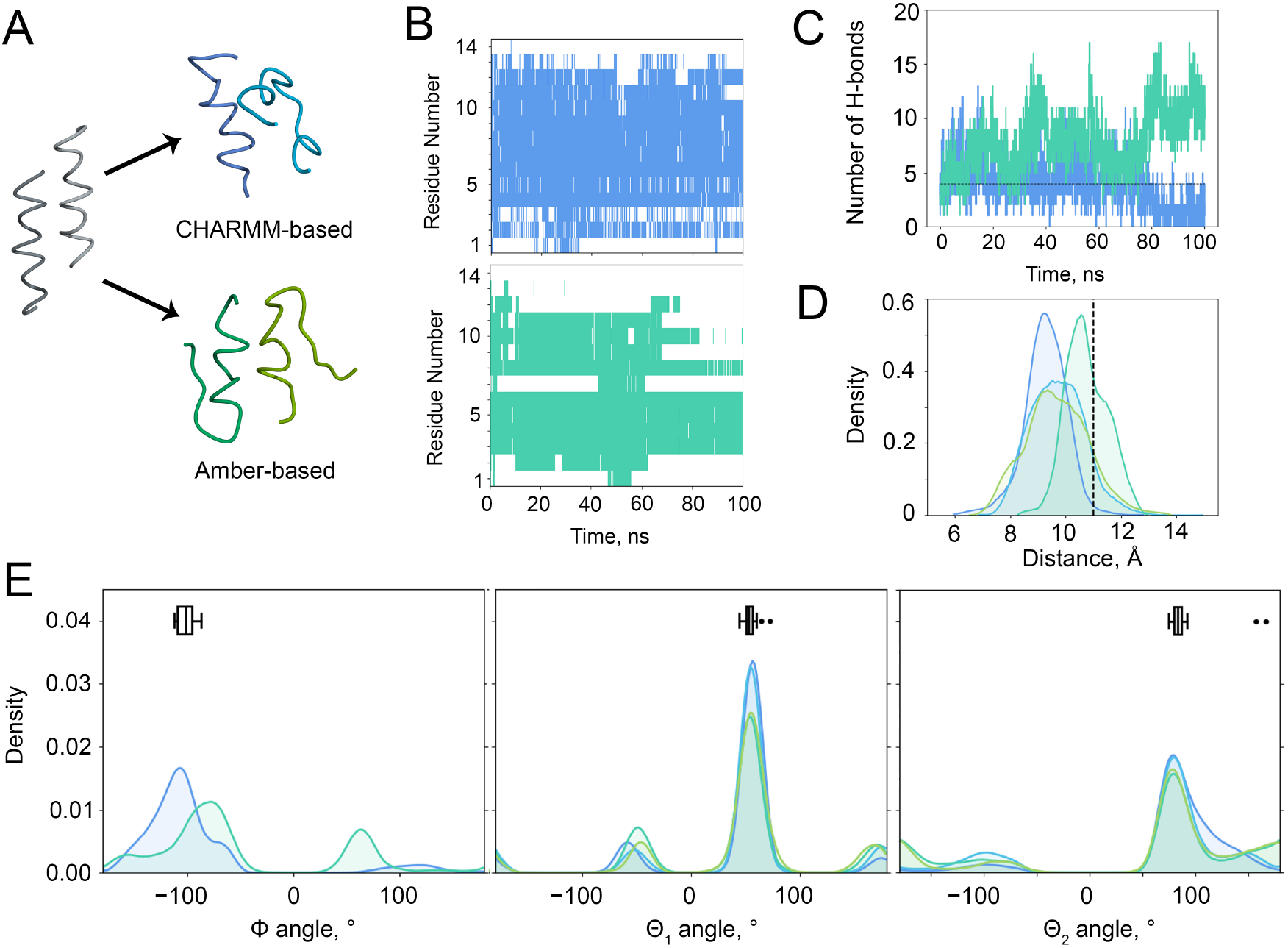
A. Last conformations sampled from the crystallographic structure of H5 for both CHARMM-(blue) and Amber-based forcefields (green). B. Time evolution of the helicity by residue. C. Number of interchains H-bonds with the number of interchain H-bonds in the crystallographic structure as reference (bold dashed line). D. Distribution of the distance of the centre of mass of both helices. E. Distribution of the backbone dihedrals Φ, Θ_1_ and Θ_2_ for each chain independently. For all figures, the crystallographic structure is represented in black (bold dashed lines, whisker boxes), CHARMM-based force field in blue, and Amber-based force field in green.

The two-helix bundle of H5 exhibits destabilization during the 100 ns simulation. However, intrachain interactions within each helix are largely preserved, and the helical folding remains partially conserved with both CHARMM and Amber-based force fields (Figure 3A).

An analysis of helicity reveals distinct behaviors differing between the two force fields. With the CHARMM-based force field, the helices maintain a steady mean helical content of approximately 9 residues throughout the simulation. In contrast, with the Amber-based force field, they show a decrease in helicity, reaching half their initial value by 100 ns (Figure 3B).

Despite the reduction in helicity observed with the Amber-based force field, interchain interactions actually increase compared to those in the CHARMM-based force field (Figure 3C). For both the CHARMM-based and Amber-based force field, approximately half of the interchain interactions consist of backbone-backbone hydrogen bonds. Moreover, in the CHARMM-based force field, there are more non-specific backbone-sidechain interactions than specific sidechain-sidechain interactions, whereas the opposite trend is observed for the Amber-based force field. Furthermore, when examining the distance between the centers of mass of the chains, we observe that the system simulated with the CHARMM-based force field forms a more tightly packed homodimer compared to both the crystallographic structure and the system simulated with the Amber-based force field (Figure 3D).

The Gaussian mixture distributions of dihedral angles reveal a consistent number of peaks and sampled angles for both force fields, similar to those observed with C5. Notably, with the CHARMM-based force field, the simulation more frequently samples the dihedral angles defined in the crystallographic structure, reinforcing its alignment with experimental data (Figure 3E).

### 2.3 Crystal patches

As a next step, we will next assess the effect of system size by analyzing crystal patches of both C5 and H5 helices using the two force fields. This approach will allow us to evaluate how well each force field reproduces the collective dynamics and assembly properties of more complex oligourea systems. To ensure computational accessibility while maintaining biotechnological relevance, we carefully selected the sizes of the crystal patches for our simulations. For helix C5, which does not naturally assemble into a nanopore, we constructed a patch composed of 58 helices to accurately represent the global crystal structure (Figure 4A).

**Figure 4:**
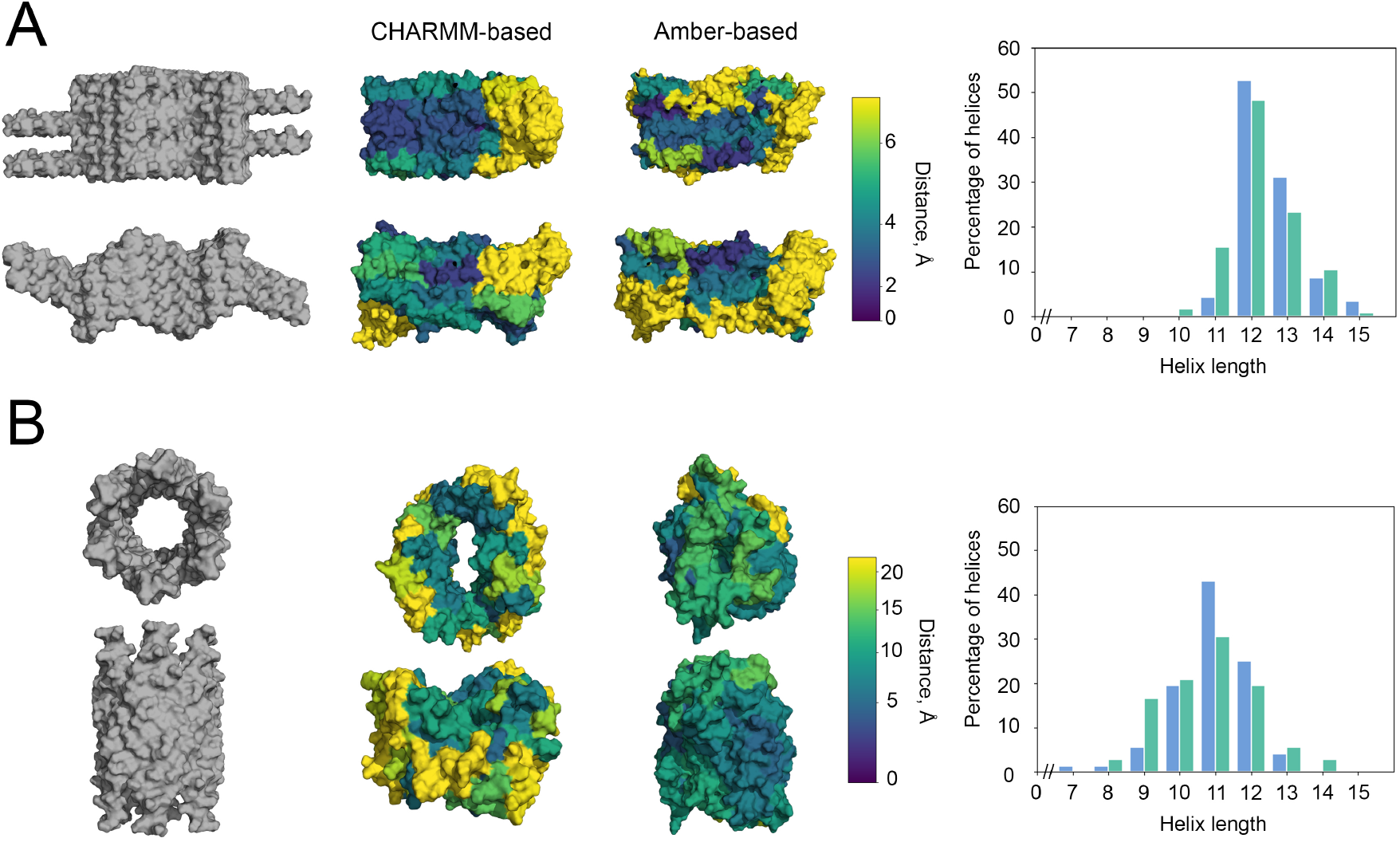
Small patches of both C5 (A) and H5 helices (B). Left. Initial structure of the crystal representing the brick-like patch of C5 helices and the nanopore formed by H5. Middle. Deviation of each constituting helix from its initial conformation. The color gradient is normalized by a power law with *V*_*min*_ = 0 and *V*_*max*_ is the maximum value of the 3rd quartile between two force fields. *γ* = 1.5 (C5) or 0.7 (H5). Right. Mean helix length.

For helix H5, we initially designed a single nanopore with a height compatible with an average lipid bilayer height of 40 Å. This nanopore is an assembly of 36 helices, arranged in a superhelical structure that replicates the twisting geometry observed in the helical transmembrane domains of protein channels. This configuration enables the study of self-assembly and functional dynamics of H5 in a relevant context. (Figure 4B). In a second step, we created a larger system formed by multiple nanopores. This system consists of seven nanopores, assembled in a cylindrical honeycomb-like structure, providing a more realistic representation of a functional oligourea-based membrane pore architecture (Figure 6). To accelerate the conformational sampling of the crystal patches, we performed two simulation replicas for 100 ns at 300 K, followed by an additional 100 ns at 330 K. As both replicas show similar behaviour, we chose to describe in detail only one of them.

#### 2.3.1 Small crystal patches

##### Patch of C5 helices

Under these conditions, the C5 foldamer assembles into a brick-like structure, which retains its global shape in simulations with either CHARMM-based or Amber-based force fields (Figure 4A, left). However, the four spanning helices (those in direct contact with the bulk of the structure) deviate noticeably from their initial conformation.

When assessing folding and spatial changes in helix orientation (measured as the deviation from the initial crystallographic structure) we found that conformational deviations increase with distance from the center of the structure, regardless of the force field used (Figure 4A, middle). This trend is primarily driven by solvent exposure, though the distribution of these changes is not radially symmetric. Instead, specific regions (particularly those more exposed to the solvent) undergo greater conformational shifts than others. For the system simulated with the CHARMM-based force field, conformational changes are predominantly localized within the spanning helices of the crystallographic patch, which are the most solvent-exposed regions. Specifically, more than 75% of helices exhibit a deviation of less than 5 Å, with 50% deviating by 3.7 Å or less. In contrast, for the system simulated with the Amber-based force field, the C5 helix patch displays conformational deviations not only in the spanning helices but also in additional solvent-exposed regions of the crystal patch. Among all helices, 75% deviate by less than 7 Å, and 50% deviate by less than 4 Å. Overall, the CHARMM-based force field results in less deviation from the initial crystal conformation compared to the Amber-based force field, indicating greater structural stability.

Beyond spatial variations, these changes also have structural consequences, as evidenced by a reduction in the number of helical residues for both force fields (Figure 4A, right). The distribution of mean helicity follows a Gaussian pattern in both cases, peaking at an average helix length of 12 residues. Notably, the CHARMM-based force field demonstrates a slightly better preservation of helicity, with 95% of helices maintaining a length above 12 residues, compared to 85% with the Amber-based force field.

##### Patch of H5 helices

When simulating the pore composed of H5 helices, the cylindrical shape is preserved in both force fields. However, the CHARMM-based force field results in a slightly reduced height of the pore, though it remains compatible with membrane embedding.

As observed with the C5 helix crystal patch, we analyzed the deviation from the original crystallographic structure of the pore (Figure 4B). Contrary to previous findings with the C5 crystal patch, the system simulated with the CHARMM-based force field exhibits greater internal conformational changes, with 75% of helices showing deviations of 22 Å or less. In comparison, the system simulated with the Amber-based force field demonstrates less deviation, with 75% of helices deviating by 12 Å or less (Figure 4B, middle). Despite the greater variation of individual helices, the CHARMM-based system maintains a large, open, oval-shaped central pore. Interestingly with the CHARMM-based force field, this does not correspond to a significant loss of helical folding (Figure 4B, right). The mean helicity distribution remains Gaussian, with a peak at an average helix length of 11 residues. This peak represents 45% of helices with the CHARMM-based force field and 30% with the Amber-based. Overall, the CHARMM-based force field demonstrates better preservation of helical folding, with 73% of helices maintaining a length above 11 residues, compared to 56% in the Amber-based force field.

When focusing on the pore formed by H5 helices, we analyzed the time evolution of the average volume of cavities with volumes greater than 1000 Å^3^ (Figure 5A). With both force fields, the pore shows a gradual decrease in cavity volume, reaching up to four times lower than the volume observed in the crystallographic structure. With the CHARMM-based force field, the decrease rate is higher and then increases, surpassing the cavity volume observed with the Amber-based force field. In contrast, the system simulated with the Amber-based force field exhibits a linear decrease in cavity volume over time (mean values averaged over two replicates).

**Figure 5:**
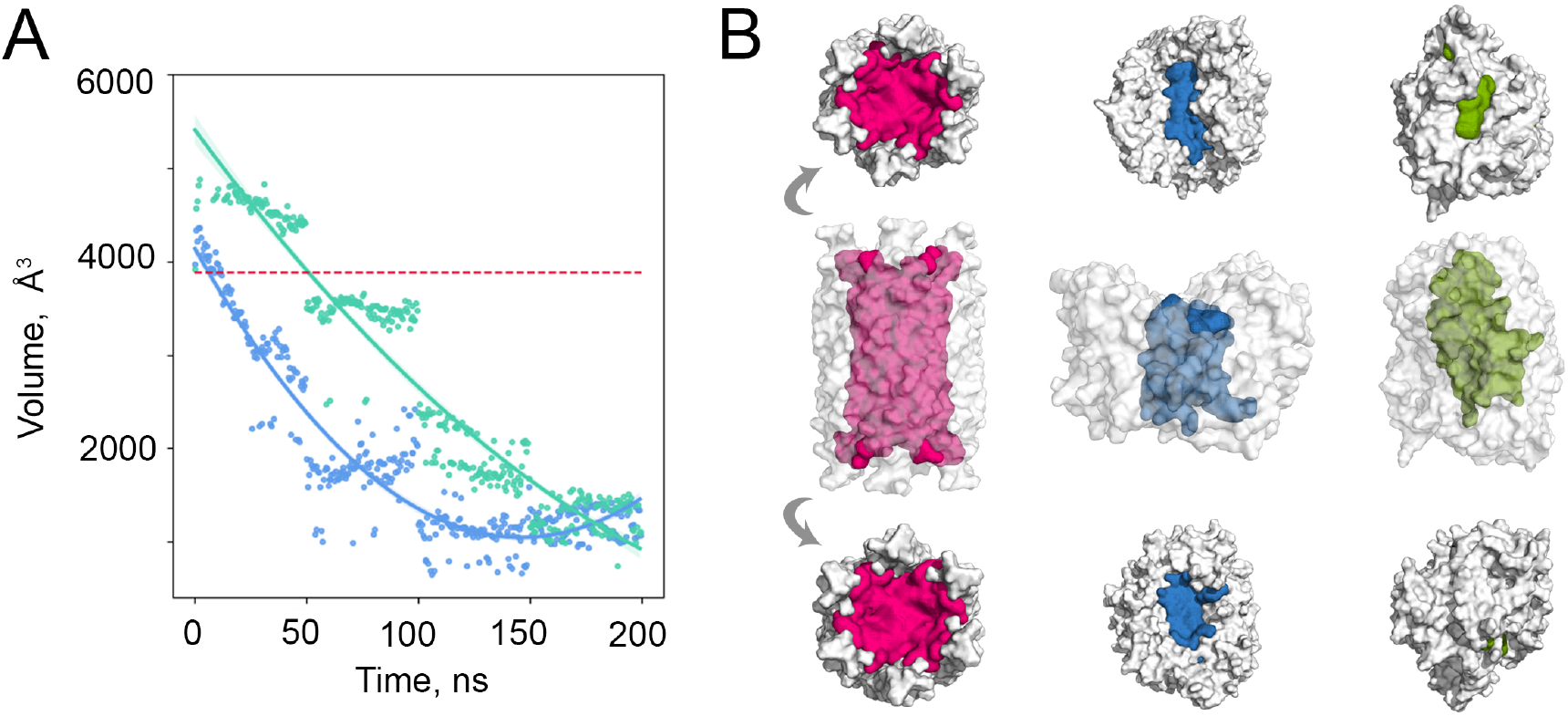
A. Time evolution of the main pore volume. B. Pore shape of the crystal nanopore and the last sampled conformation. The CHARMM-based force field is in blue and the Amber-based in green. The initial pore location and reference value are represented in bold dashed pink.

At the final sampled conformation, the pore entries in both force fields exhibit noticeable narrowing (Figure 5B). However, a key distinction emerges between the two force fields. In the CHARMM-based simulation, the pore remains open on both sides, maintaining a continuous central cavity with a volume of 10800 Å^3^ (Figure 5B, middle). In contrast, the Amber-based simulation results show a pore that is completely closed on one side and severely constricted on the other, with a cavity volume of 12900 Å^3^ (Figure 5B, right).

This analysis highlights differences in pore stability between the two force fields, with the CHARMM-based force field preserving a more accessible pore geometry compared to the system simulated with the Amber-based force field.

### 2.4 Honeycomb patch of H5

While these findings are promising for selecting the most appropriate force field for oligourea modeling, it is important to note that, under crystallographic conditions, oligourea pores typically exist as larger, interconnected assemblies. To better reflect this experimental context, we will now discuss the stability of a larger system composed of seven nanopores (Figure 6).

**Figure 6:**
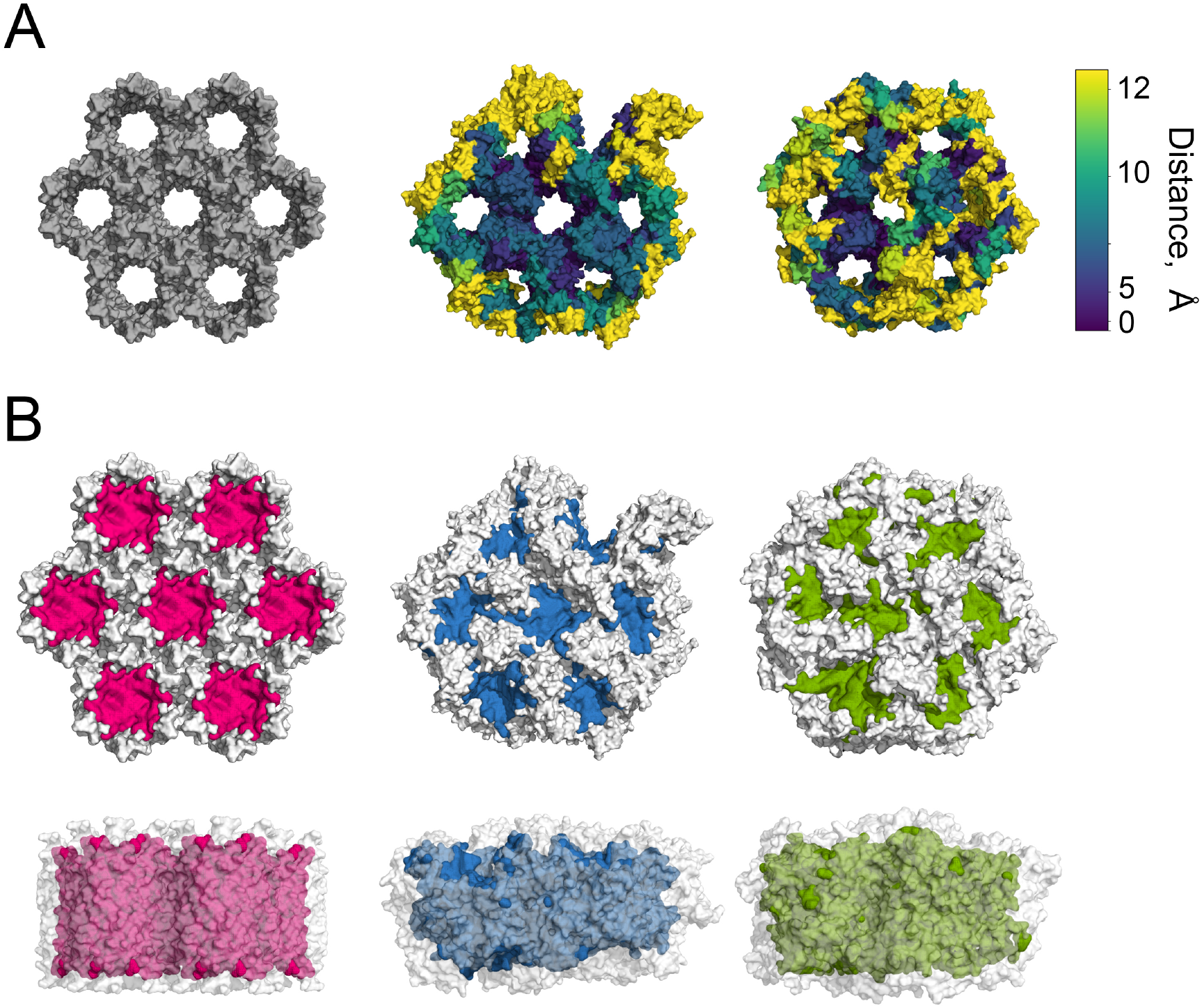
A. Initial structure of the honeycomb crystal structure (left) and deviation of each constituting helix from its initial conformation. The color gradient is normalized by a power law with *V*_*min*_ = 0 and *V*_*max*_ is the maximum value of the 3rd quartile between two force fields. *γ* = 2. B. Pore shape and location from the top and side view. The initial pore locations are represented in pink.

Compared to the regular honeycomb shape of the initial crystallographic structure, the simulated systems using different force fields exhibit distinct patterns of structural degradation, yet both retain an overall conservation of the original shape. Similar to the small nanopore, the honeycomb structure displays the most pronounced conformational and spatial changes in the helices located at the solvent interface. Notably, the system simulated with the middle pore in the system simulated with the CHARMM-based force field demonstrates greater internal stability compared to the system simulated with the Amber-based force field (Figure 6A).

Regarding porosity, the honeycomb structure largely retains its porous nature. However, in the system simulated with the CHARMM-based force field, a pore in contact with the solvent opened. In contrast, the system simulated with the Amber-based force field maintains a closed shape at the solvent interface.

Additionally, the pores in the Amber-based system appear slightly looser compared to both the initial crystallographic structure and the CHARMM-based system. These observations are further supported by the visualization of the pore cavities (Figure 6B).

Focusing on the “protected” middle pore, the mean helicity distribution for the systems simulated with both CHARMM- and Amber-based force fields follow a Gaussian-shaped profile, with a peak at a helix length of 11 residues, consistent with observations in the single-pore system. However, a clear distinction emerges in the mean helical content between the two force fields: the system simulated with the CHARMM-based force field maintains over 65% of helices at a length of 11 residues, with more than 75% of helices exceeding this length. In contrast, the Amber-based force field samples only 35% of helices at a length of 11 residues, with 60% of helices exceeding this threshold (Figure 7A).

**Figure 7:**
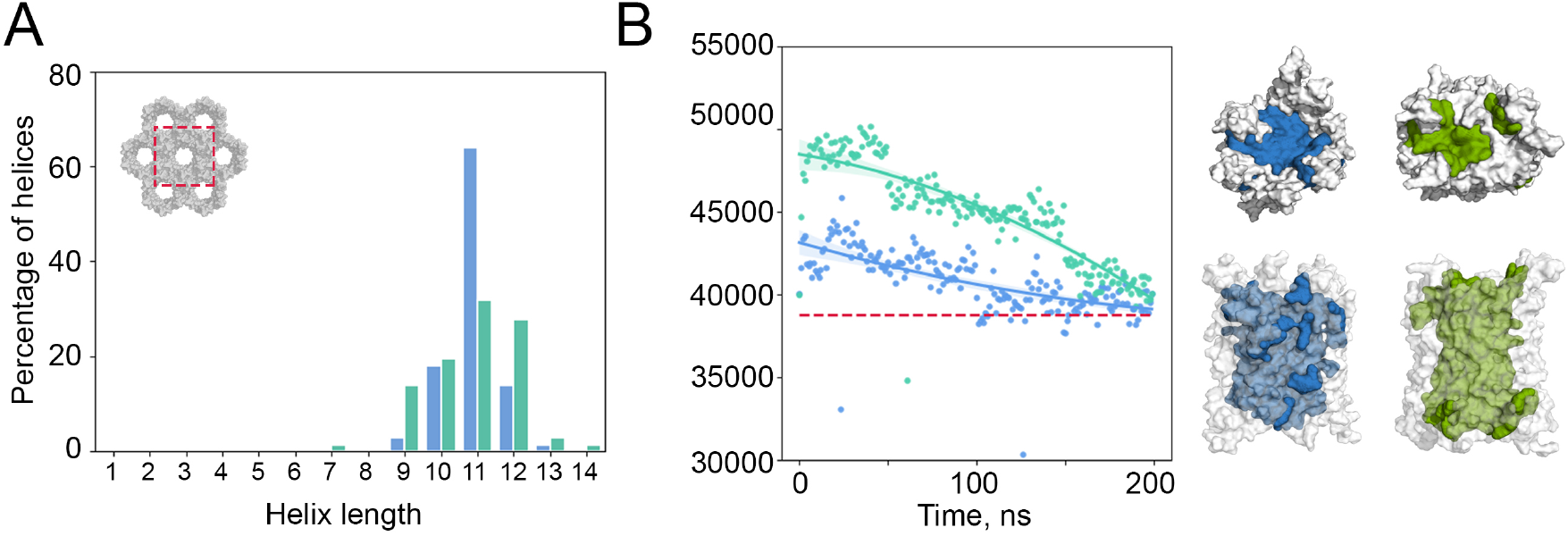
A. Mean helix length. B. Time evolution of the main pore volume (left) and pore shape of the last sampled conformation. The CHARMM-based force field is in blue and the Amber-based one in green. The initial pore location and volume reference value are represented in bold dashed pink.

When examining the time evolution of pore volume, we observe a fast initial increase in volume during the first few nanoseconds, followed by a gradual decrease with similar rate of the two systems, until the volume stabilizes near that of the crystallographic structure for both CHARMM- and Amber-based force field (Figure 7B, left).

Despite differences in helicity, the middle nanopore retains its overall shape even with partially unfolded helices. In both cases, the global shape of the middle pore remains conserved, demonstrating that the macroscopic architecture is preserved regardless of the force field used (Figure 7B, right). We nevertheless note that in the last sampled conformation of the system simulated with the Amber-based force field, one of the extremities of the nanopore begins to be “covered” by one of the composing helices. We could naturally expect that in a larger simulation timescale, this opening would be at least partially closed.

## 3 Conclusion

In this study, we aimed to develop an accurate force field for the *in silico* study of oligourea foldamers. Given the structural closeness of oligourea foldamers to proteins, we chose to adapt parameters from two well-established protein force field families: CHARMM and Amber.

Using available crystallographic structures of oligourea foldamers, we compared molecular dynamics (MD) simulation ensembles, focusing on key structural features such as folding patterns, B-factors, and pore shapes. It is important to clarify that our objective was not to determine which oligourea foldamer is better suited for water conduction, but rather to select a force field that reliably maintains structural integrity.

We began by adjusting the partial charges and atom types of oligourea residues, with a particular focus on backbone atoms. For the CHARMM-based force field, we assigned charges and atom types based on the CHARMM36m protein force field. For the Amber-based force field, charges and atom types were assigned using the GAFF2 force field and the AM1-BCC charge model with an additional charge correction of the C*α*.

Starting from these force fields, we evaluated two oligourea foldamers: C5 (aliphatic) and H5 (amphiphilic), using increasing system sizes and comparing them to experimental structural features. MD simulations of these systems with our CHARMM- and Amber-based force fields revealed that, regardless of the solvent, C5 exhibits consistent features with either force field. For helix C5, both in a single helix and an extended crystal patch, the CHARMM-based force field better reproduces experimental features. For H5, we reached the same conclusion, with additional insights into pore shape: in a single nanopore, the CHARMM-based force field maintains the pore shape, opening, and helicity, whereas the Amber-based force field tends to close the pore on one side. In a further extended honeycomb-shaped system, both force fields result in similar pore size and shape, but the CHARMM-based force field better preserves helicity in the “protected” central pore.

Globally, the CHARMM-based force field demonstrates superior performance in preserving the structural integrity of C5 and H5 helix assemblies, whether in a single helix or a larger, extended crystal configuration. This consistency highlights its reliability for modeling oligourea-based systems, making it our preferred choice for future studies.

However, our models may have limitations, as we did not yet replicate the conditions employed in water desalination processes, such as being embedded in a membrane. Building on these results and our force field selection, we are now simulating the H5 multipore system in a membrane environment to accurately reproduce the global environment of this oligourea when embedded in a membrane. This future study will focus first on evaluating the structural and dynamical features of the system at osmotic equilibrium. Subsequently, we will assess its stability under reverse osmosis conditions, including high-pressure and osmotic gradients of salt, using a double membrane system.

## 4 Materials and Methods

### 4.1 Force field construction

#### CHARMM-based force field

We assigned the same charges and atom types as those defined for the protein force field CHARMM36m for all common atoms, including the backbone and side chains. The CH_2_ atom types and charges were assigned according to the CHARMM36m force field [22], while the backbone N’H’ of the urea group was assigned types and charges identical to those of a peptide backbone amide group.

#### Amber-based force field

First, we generated the topology of a tri-mer (Boc-uG-uX-uG-NMe) for each residue X using the CHARMM-GUI Ligand Reader module [29]. We then used the Membrane Builder module to produce GROMACS-formatted topologies and parameters for GAFF2 force fields with AM1-BCC charges [23, 24], including additional parameters for non-bonded interactions to ensure compatibility with potential membrane simulations. From these outputs, we extracted the individual parameter blocks for each oligourea residue.

This force field presented two main challenges. The first was the discrepancy between the charges of the C*α* atom in the latest Amber protein force field (ff19SB) and the generated GAFF2 parameters. Additionally, the total charge of each residue did not match the expected values (0 for neutral residues, -1 or +1 for charged residues such as uD/uE and uK/uR). To resolve this mismatch, we retained the charges of all atoms and adjusted the C*α* charges to ensure the total charge of each residue was as required.

### 4.2 Preparation of the systems

#### Crystallographic structures

The crystallographic structure of the oligoureas were obtained from the *Cambridge Crystallographic Data Centre* (CCDC), with the following accession numbers: 836812 for helix C5 and 1030454 for helix H5.

#### Crystal patches

Samples of the C5 and H5 crystals were constructed using VESTA 3.5.8 [30] by expanding the boundaries of the unit cell. For the C5 patch and H5 nanopore, the unit-cell boundaries were set to 0–2 along the x and y axes and 0–1.5 along the z axis. For the honeycomb H5 multipore assembly, the unit-cell boundaries were extended to 0–4 along the x and y axes and 0–1 along the z axis.

The structures with added hydrogens were then placed in either an octahedral box (for single-helix systems) or a cubic box (for crystal patches), ensuring a minimum distance of 1.2 nm between the oligourea and the box boundary. The systems were subsequently solvated in TIP3P water with a 0.15 mM NaCl concentration, or in pure methanol (for the C5 helix only).

### 4.3 MD simulations

All-atom molecular dynamics (MD) simulations of the single-helix and crystal patches of compound C5 and helix H5 were performed using GROMACS 2023.2 [31]. To ensure consistency and enable direct comparison between the CHARMM-based and Amber-based force fields, the equilibration and simulation procedures were kept identical for all systems.

#### Equilibration

Before simulation, the systems were relaxed following the protocol. First, 10,000 steps of steepest descent minimization were performed with a harmonic force constraint of 4000 kJ/mol applied to the oligourea to prevent structural distortions. With the same force constraint, a 500 ps NPT equilibration was conducted to remove artificial voids within the simulation box, followed by a 500 ps NVT equilibration to adjust the box density. Finally, a 500 ps NPT equilibration was performed with gradually decreasing force constraints (2000, 1000, 500, 200, and 50 kJ/mol) to allow the slow and controlled relaxation of the oligourea. For all equilibration steps, the V-rescale thermostat (300 K) [32] and C-rescale barostat [33] (1 bar, isotropic coupling) were used to maintain temperature and pressure stability.

#### Production

Duplicates of the single-helix systems of C5 and helix H5 were simulated for 100 ns at 300 K and 1 bar, using the V-rescale thermostat and C-rescale barostat with isotropic coupling [32, 33]. Under the same conditions, the crystal patches were simulated for 100 ns at 300 K, followed by an additional 100 ns at 330 K. For all simulations, the short-range non-bonded interactions were truncated at a cutoff of 1.2 nm, and the long-range electrostatic interactions were evaluated using the Particle Mesh Ewald (PME) method [34]. Bonds involving hydrogen atoms and their heavy atoms were constrained using the LINCS algorithm [35]. The equations of motion were integrated with the leap-frog algorithm [36] with a time step of 2 fs.

### 4.4 Analysis

As an initial step in trajectory processing, we removed rotational and translational motions by aligning all structures to the first frame using their backbone atoms (N, C*α*, C’, N’, C, O).

Unless otherwise specified, all analyses were conducted using the CPPTRAJ module (version 6.24.0) from AmberTools 24.8 [37].

#### Hydrogen Bonds (H-bonds)

H-bonds were identified using geometric criteria based on donor and acceptor (A) atoms (N and O). An H-bond was defined if the distance DA *≤* 3.6 Å and the angle DHA *≥* 110°. Only H-bonds with a frequency *≥* 0.01 were retained for further analysis.

#### Helicity

Following the H-bond analysis, a residue i was considered part of a helix if an H-bond existed between the backbone of residues *i*/*i −* 2 and *i*/*i* + 2.

#### Backbone Dihedrals

Backbone dihedral angles were calculated based on the crystallographic definitions: Φ (C–N–C*α*–C’), Θ_1_ (N–C*α*–C’–N’), Θ_2_ (C*α*–C’–N’–C).

#### Backbone Fluctuations

Backbone fluctuations were calculated using the average conformation after fitting to the backbone atoms.

#### Pore cavities

Pore cavities were computed using CavitOmiX 1.0 [38] with a probe radius of 1.4 Å, including hydrogen atoms. For clarity, only cavities with a volume V*≥*1000 Å^3^ were considered in the analysis.

## Acknowledgements

We thank the French National Agency for Research for funding (Grant Aquafoldamers ANR-23-CE06-0030,LABEX DYNAMO ANR-LABX-011, EQUIPEX CACSICE ANR-11-EQPX-0008). Molecular simulations were performed using HPC computing resources from GENCI-/TGCC/CINES/IDRIS (Grant A0190701714 to MB). M.B. thanks Sesame Ile-de-France.

## 5 Supplementary Material

**Table 1:**
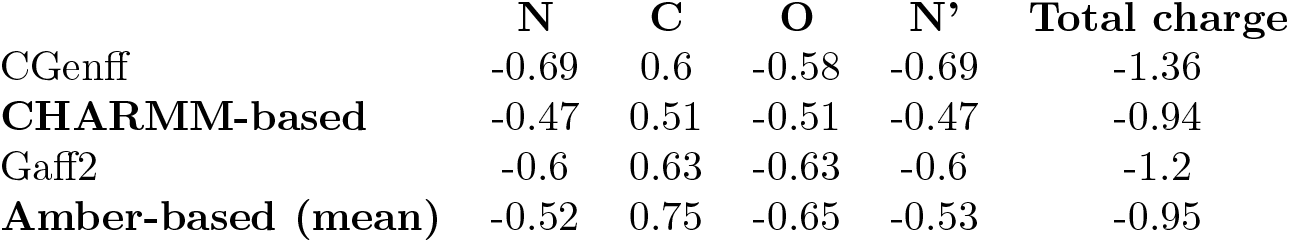
Comparison of an oligourea urea group charges with the urea molecule (NH_2_-(C=O)-NH_2_) for our analogy-based forcefields and the general purose force fields Cgenff and Gaff2 with AM1-BCC charges.

|  | N | C | O | N' | Total charge |
| --- | --- | --- | --- | --- | --- |
| CGenff | -0.69 | 0.6 | -0.58 | -0.69 | -1.36 |
| **CHARMM-based** | -0.47 | 0.51 | -0.51 | -0.47 | -0.94 |
| Gaff2 | -0.6 | 0.63 | -0.63 | -0.6 | -1.2 |
| **Amber-based (mean)** | -0.52 | 0.75 | -0.65 | -0.53 | -0.95 |

**Table 2:** Comparison of an oligourea urea group charges with the 1,3-dimethylurea molecule (CH_3_-NH-(C=O)-NH-CH_3_) with our analogy-based force fields and the general purpose force fields Cgenff and Gaff2 with AM1-BCC charges.

|  | N | C | O | N' | Total charge |
| --- | --- | --- | --- | --- | --- |
| CGenff | -0.44 | 0.25 | -0.41 | -0.44 | -1.04 |
| **CHARMM-based** | -0.47 | 0.51 | -0.51 | -0.47 | -0.94 |
| Gaff2 | -0.53 | 0.65 | -0.64 | -0.53 | -1.05 |
| **Amber-based (mean)** | -0.52 | 0.75 | -0.65 | -0.53 | -0.95 |

## References

[1] Water – at the Center of the Climate Crisis. United Nations. 2025. url: https://www.un.org/en/climatechange/science/climate-issues/water.

[2] Auindrila Biswas et al. “Water scarcity: A global hindrance to sustainable development and agricultural production – A critical review of the impacts and adaptation strategies”. In: Cambridge Prisms: Water 3 (2025), e4. issn: 2755-1776. doi: 10.1017/wat.2024.16. URL: https://www.cambridge.org/core/product/identifier/S2755177624000169/type/journal_article (visited on 12/25/2025).

[3] Junguo Liu et al. “Water scarcity assessments in the past, present and future”. In: Earth’s Future 5.6 (June 2017), pp. 545–559. ISSN: 2328-4277. doi: 10.1002/2016EF000518.

[4] Ahmed Alkaisi, Ruth Mossad, and Ahmad Sharifian-Barforoush. “A Review of the Water Desalination Systems Integrated with Renewable Energy”. In: Energy Procedia 110 (Mar. 2017), pp. 268–274. ISSN: 18766102. doi: 10.1016/j.egypro.2017.03.138. URL: https://linkinghub.elsevier.com/retrieve/pii/S1876610217301686 (visited on 12/18/2025).

[5] Muhammad Qasim et al. “Reverse osmosis desalination: A state-of-the-art review”. In: Desalination 459 (June 2019), pp. 59–104. ISSN: 00119164. doi: 10.1016/j.desal.2019.02.008. URL: https://linkinghub.elsevier.com/retrieve/pii/S0011916418325037 (visited on 12/18/2025).

[6] Yousef A. Tayeh. “A comprehensive review of reverse osmosis desalination: Technology, water sources, membrane processes, fouling, and cleaning”. In: Desalination and Water Treatment 320 (Oct. 2024), p. 100882. ISSN: 19443986. doi: 10.1016/j.dwt.2024.100882. URL: https://linkinghub.elsevier.com/retrieve/pii/S1944398624203921 (visited on 12/18/2025).

[7] Frédéric H. Login and Lene N. Nejsum. “Aquaporin water channels: roles beyond renal water handling”. In: Nature Reviews Nephrology 19.9 (Sept. 2023), pp. 604–618. ISSN: 1759-5061, 1759-507X. doi: 10.1038/s41581-023-00734-9. URL: https://www.nature.com/articles/s41581-023-00734-9 (visited on 12/18/2025).

[8] Abouzar Azarafza et al. “Aquaporin-Based Biomimetic Membranes for Low Energy Water Desalination and Separation Applications”. In: Advanced Functional Materials 33.21 (May 2023), p. 2213326. ISSN: 1616-301X, 1616-3028. doi: 10.1002/adfm.202213326. URL: https://advanced.onlinelibrary.wiley.com/doi/10.1002/adfm.202213326 (visited on 12/18/2025).

[9] Jie Shen et al. “Artificial channels for confined mass transport at the sub-nanometre scale”. In: Nature Reviews Materials 6.4 (Jan. 21, 2021), pp. 294–312. ISSN: 2058-8437. doi: 10.1038/s41578-020-00268-7. URL: https://www.nature.com/articles/s41578-020-00268-7 (visited on 12/18/2025).

[10] Chiranjit Dutta et al. “Nature-inspired synthetic oligourea foldamer channels allow water transport with high salt rejection”. In: Chem 9.8 (Aug. 2023), pp. 2237–2254. ISSN: 24519294. doi: 10.1016/j.chempr.2023.04.007. URL: https://linkinghub.elsevier.com/retrieve/pii/S2451929423001870 (visited on 12/25/2025).

[11] Piotr Zielenkiewicz, Danuta Plochocka, and Andrzej Rabczenko. “The formation of protein secondary structure”. In: Biophysical Chemistry 31.1 (Aug. 1988), pp. 139–142. ISSN: 03014622. doi: 10.1016/0301-4622(88)80018-8. URL: https://linkinghub.elsevier.com/retrieve/pii/0301462288800188 (visited on 12/25/2025).

[12] Samuel H. Gellman. “Foldamers: A Manifesto”. In: Accounts of Chemical Research 31.4 (Apr. 1, 1998), pp. 173–180. ISSN: 0001-4842, 1520-4898. doi: 10.1021/ar960298r. URL: https://pubs.acs.org/doi/10.1021/ar960298r (visited on 12/25/2025).

[13] Saquib Farooq et al. “Rapid Water Permeation by Aramid Foldamer Nanochannels With Hydrophobic Interiors”. In: Angewandte Chemie International Edition 64.22 (May 26, 2025), e202504170. ISSN: 1433-7851, 1521–3773. doi: 10.1002/anie.202504170. URL: https://onlinelibrary.wiley.com/doi/10.1002/anie.202504170 (visited on 12/25/2025).

[14] Arundhati Roy et al. “Foldamer-based ultrapermeable and highly selective artificial water channels that exclude protons”. In: Nature Nanotechnology 16.8 (Aug. 2021), pp. 911–917. ISSN: 1748-3387, 1748-3395. doi: 10.1038/s41565-021-00915-2. URL: https://www.nature.com/articles/s41565-021-00915-2 (visited on 12/25/2025).

[15] W. Seth Horne and Samuel H. Gellman. “Foldamers with Heterogeneous Backbones”. In: Accounts of Chemical Research 41.10 (Oct. 21, 2008), pp. 1399–1408. ISSN: 0001-4842, 1520-4898. doi: 10.1021/ar800009n. URL: https://pubs.acs.org/doi/10.1021/ar800009n (visited on 12/25/2025).

[16] Karl T. Debiec et al. “Further along the Road Less Traveled: AMBER ff15ipq, an Original Protein Force Field Built on a Self-Consistent Physical Model”. In: Journal of Chemical Theory and Computation 12.8 (Aug. 9, 2016), pp. 3926–3947. ISSN: 1549-9618, 1549-9626. doi: 10.1021/acs.jctc.6b00567. URL: https://pubs.acs.org/doi/10.1021/acs.jctc.6b00567 (visited on 12/15/2025).

[17] András Wacha, Zoltán Varga, and Tamás Beke-Somfai. “Comparative Study of Molecular Mechanics Force Fields for-Peptidic Foldamers: Folding and Self-Association”. In: Journal of Chemical Information and Modeling 63.12 (June 26, 2023), pp. 3799–3813. ISSN: 1549-9596, 1549-960X. doi: 10.1021/acs.jcim.3c00175. URL: https://pubs.acs.org/doi/10.1021/acs.jcim.3c00175 (visited on 11/07/2025).

[18] George A. Khoury et al. “Forcefield NCAA: *Ab Initio* Charge Parameters to Aid in the Discovery and Design of Therapeutic Proteins and Peptides with Unnatural Amino Acids and Their Application to Complement Inhibitors of the Compstatin Family”. In: ACS Synthetic Biology 3.12 (Dec. 19, 2014), pp. 855–869. ISSN: 2161-5063, 2161-5063. doi: 10.1021/sb400168u. URL: https://pubs.acs.org/doi/10.1021/sb400168u (visited on 12/15/2025).

[19] K. Vanommeslaeghe et al. “CHARMM general force field: A force field for drug-like molecules compatible with the CHARMM all-atom additive biological force fields”. In: Journal of Computational Chemistry 31.4 (Mar. 2010), pp. 671–690. ISSN: 0192-8651, 1096-987X. doi: 10.1002/jcc.21367. URL: https://onlinelibrary.wiley.com/doi/10.1002/jcc.21367 (visited on 11/07/2025).

[20] Dina T. Mirijanian et al. “Development and use of an atomistic CHARMM-based force-field for peptoid simulation”. In: Journal of Computational Chemistry 35.5 (Feb. 15, 2014), pp. 360–370. ISSN: 01928651. doi: 10.1002/jcc.23478. URL: https://onlinelibrary.wiley.com/doi/10.1002/jcc.23478 (visited on 11/07/2025).

[21] Jing Huang et al. “CHARMM36m: an improved force field for folded and intrinsically disordered proteins”. In: Nature Methods 14.1 (Jan. 2017), pp. 71–73. ISSN: 1548-7091, 1548-7105. doi: 10.1038/nmeth.4067. URL: https://www.nature.com/articles/nmeth.4067 (visited on 11/07/2025).

[22] Chuan Tian et al. “ff19SB: Amino-Acid-Specific Protein Backbone Parameters Trained against Quantum Mechanics Energy Surfaces in Solution”. In: Journal of Chemical Theory and Computation 16.1 (Jan. 14, 2020), pp. 528–552. ISSN: 1549-9618, 1549-9626. doi: 10.1021/acs.jctc.9b00591. URL: https://pubs.acs.org/doi/10.1021/acs.jctc.9b00591 (visited on 12/16/2025).

[23] Xibing He et al. “A fast and high-quality charge model for the next generation general AMBER force field”. In: The Journal of Chemical Physics 153.11 (Sept. 21, 2020), p. 114502. ISSN: 0021-9606, 1089-7690. doi: 10.1063/5.0019056. URL: https://pubs.aip.org/jcp/article/153/11/114502/199591/A-fast-and-high-quality-charge-model-for-the-next (visited on 12/18/2025).

[24] Junmei Wang et al. “Development and testing of a general amber force field”. In: Journal of Computational Chemistry 25.9 (July 15, 2004), pp. 1157–1174. ISSN: 0192-8651, 1096-987X. doi: 10.1002/jcc.20035. URL: https://onlinelibrary.wiley.com/doi/10.1002/jcc.20035 (visited on 12/18/2025).

[25] Lucile Fischer et al. “The Canonical Helix of Urea Oligomers at Atomic Resolution: Insights Into Folding-Induced Axial Organization”. In: Angewandte Chemie International Edition 49.6 (Feb. 2010), pp. 1067–1070. ISSN: 1433-7851, 1521-3773. doi: 10.1002/anie.200905592. URL: https://onlinelibrary.wiley.com/doi/10.1002/anie.200905592 (visited on 11/07/2025).

[26] Peter J. Bond et al. “Membrane protein dynamics and detergent interactions within a crystal: A simulation study of OmpA”. In: Proceedings of the National Academy of Sciences 103.25 (June 20, 2006), pp. 9518–9523. ISSN: 0027-8424, 1091-6490. doi: 10.1073/pnas.0600398103. URL: https://pnas.org/doi/full/10.1073/pnas.0600398103 (visited on 11/07/2025).

[27] Gavin W. Collie et al. “Shaping quaternary assemblies of water-soluble non-peptide helical foldamers by sequence manipulation”. In: Nature Chemistry 7.11 (Nov. 2015), pp. 871–878. ISSN: 1755-4330, 1755-4349. doi: 10.1038/nchem.2353. URL: https://www.nature.com/articles/nchem.2353 (visited on 11/07/2025).

[28] Juliette Fremaux et al. “Condensation Approach to Aliphatic Oligourea Foldamers: Helices with *N*-(Pyrrolidin-2-ylmethyl)ureido Junctions”. In: Angewandte Chemie International Edition 50.48 (Nov. 25, 2011), pp. 11382–11385. ISSN: 1433-7851, 1521-3773. doi: 10.1002/anie.201105416. URL: https://onlinelibrary.wiley.com/doi/10.1002/anie.201105416 (visited on 11/07/2025).

[29] Sunhwan Jo et al. “CHARMM-GUI: A web-based graphical user interface for CHARMM”. In: Journal of Computational Chemistry 29.11 (Aug. 2008), pp. 1859–1865. ISSN: 0192-8651, 1096-987X. doi: 10.1002/jcc.20945. URL: https://onlinelibrary.wiley.com/doi/10.1002/jcc.20945 (visited on 12/25/2025).

[30] Koichi Momma and Fujio Izumi. “*VESTA 3* for three-dimensional visualization of crystal, volumetric and morphology data”. In: Journal of Applied Crystallography 44.6 (Dec. 1, 2011), pp. 1272–1276. ISSN: 0021-8898. doi: 10.1107/S0021889811038970. URL: https://journals.iucr.org/paper?S0021889811038970 (visited on 12/25/2025).

[31] Mark James Abraham et al. “GROMACS: High performance molecular simulations through multi-level parallelism from laptops to supercomputers”. In: SoftwareX 1-2 (Sept. 2015), pp. 19–25. ISSN: 23527110. doi: 10.1016/j.softx.2015.06.001. URL: https://linkinghub.elsevier.com/retrieve/pii/S2352711015000059 (visited on 12/25/2025).

[32] Giovanni Bussi, Davide Donadio, and Michele Parrinello. “Canonical sampling through velocity rescaling”. In: The Journal of Chemical Physics 126.1 (Jan. 7, 2007), p. 014101. ISSN: 0021-9606, 1089-7690. doi: 10.1063/1.2408420. URL: https://pubs.aip.org/jcp/article/126/1/014101/186581/Canonical-sampling-through-velocity-rescaling (visited on 12/25/2025).

[33] Mattia Bernetti and Giovanni Bussi. “Pressure control using stochastic cell rescaling”. In: The Journal of Chemical Physics 153.11 (Sept. 21, 2020), p. 114107. ISSN: 0021-9606, 1089-7690. doi: 10.1063/5.0020514. URL: https://pubs.aip.org/jcp/article/153/11/114107/199610/Pressure-control-using-stochastic-cell-rescaling (visited on 12/25/2025).

[34] Celeste Sagui, Lee G. Pedersen, and Thomas A. Darden. “Towards an accurate representation of electrostatics in classical force fields: Efficient implementation of multipolar interactions in biomolecular simulations”. In: The Journal of Chemical Physics 120.1 (Jan. 1, 2004), pp. 73–87. ISSN: 0021-9606, 1089-7690. doi: 10.1063/1.1630791. URL: https://pubs.aip.org/jcp/article/120/1/73/534195/Towards-an-accurate-representation-of (visited on 12/25/2025).

[35] Berk Hess et al. “LINCS: A linear constraint solver for molecular simulations”. In: Journal of Computational Chemistry 18.12 (Sept. 1997), pp. 1463–1472. ISSN: 0192-8651, 1096-987X. doi: 10.1002/(SICI)1096-987X(199709)18:12<1463::AID-JCC4>3.0.CO;2-H. URL: https://onlinelibrary.wiley.com/doi/10.1002/(SICI)1096-987X(199709)18:12%3C1463::AID-JCC4%3E3.0.CO;2-H (visited on 12/25/2025).

[36] W. F. Van Gunsteren and H. J. C. Berendsen. “A Leap-frog Algorithm for Stochastic Dynamics”. In: Molecular Simulation 1.3 (Mar. 1988), pp. 173–185. ISSN: 0892-7022, 1029-0435. doi: 10.1080/08927028808080941. URL: https://www.tandfonline.com/doi/full/10.1080/08927028808080941 (visited on 12/25/2025).

[37] David A. Case et al. “Recent Developments in Amber Biomolecular Simulations”. In: Journal of Chemical Information and Modeling 65.15 (Aug. 11, 2025), pp. 7835–7843. ISSN: 1549-9596, 1549-960X. doi: 10.1021/acs.jcim.5c01063. URL: https://pubs.acs.org/doi/10.1021/acs.jcim.5c01063 (visited on 12/25/2025).

[38] CavitOmiX. Version 1.0. 2022. URL: https://innophore.com/cavitomix.

